# Task-specific roles of local interneurons for inter -and intraglomerular signaling in the insect antennal lobe

**DOI:** 10.1101/2020.11.26.369686

**Authors:** Debora Fusca, Peter Kloppenburg

## Abstract

Local interneurons (LNs) mediate complex interactions within the antennal lobe, the primary olfactory system of insects, and the functional analog of the vertebrate olfactory bulb. In the cockroach *Periplaneta Americana*, as in other insects, several types of LNs with distinctive physiological and morphological properties can be defined. Here, we combined whole-cell patch clamp recordings and Ca^2+^ imaging of individual LNs to analyze the role of spiking and nonspiking LNs in inter- and intraglomerular signaling during olfactory information processing. Spiking GABAergic LNs reacted to odorant stimulation with a uniform rise in *[Ca*^*2+*^]_*i*_ in the ramifications of all innervated glomeruli. In contrast, in nonspiking LNs, glomerular Ca^2+^ signals were odorant specific and varied between glomeruli, resulting in distinct, glomerulus-specific tuning curves. The cell type-specific differences in Ca^2+^ dynamics support the idea that spiking LNs play a primary role in interglomerular signaling, while they assign nonspiking LNs an essential role in intraglomerular signaling.

## INTRODUCTION

Local interneurons (LNs) with markedly different functional phenotypes are crucial for odor information processing in the insect antennal lobe (AL). The AL is the first synaptic relay in the insect olfactory system, showing striking structural and functional similarities to the vertebrates’ olfactory bulb. In many regards, the LNs in the AL are the functional equivalent of granule cells, but also periglomerular and short axon cells in the vertebrate olfactory bulb (Ennis et al., 2015; Shepherd et al., 2004). They help to structure the odor representation in the AL, ultimately shaping the tuning profiles of the olfactory projection (output) neurons.

Based on initial studies, LNs originally have been characterized as GABAergic and multiglomerular (Distler, 1989; Hoskins et al., 1986; Waldrop et al., 1987). Typically, they can generate Na^+^-driven action potentials (Chou et al., 2010; Christensen et al., 1993; Husch et al., 2009a; Seki et al., 2010) or Ca^2+^-driven spikelets (Laurent and Davidowitz, 1994). Accordingly, these neurons have been associated with inhibitory interglomerular signaling, i.e., with mediating lateral inhibition to enhance contrast and to control timing and synchronization of neuronal activity (Assisi and Bazhenov, 2012; Christensen et al., 1998; Fujiwara et al., 2014; MacLeod and Laurent, 1996; Nagel and Wilson, 2016; Sachse and Galizia, 2002). Subsequent studies showed that LNs can also synthesize other potential neurotransmitters and neuromodulators (Berg et al., 2007; Chou et al., 2010; Das et al., 2011; Distler, 1990; Fusca et al., 2013, 2015; Neupert et al., 2012; Shang et al., 2007). In fact, they can be excitatory, distributing excitatory synaptic input to (projection) neurons in other glomeruli (Assisi et al., 2012; Huang et al., 2010; Yaksi and Wilson, 2010).

Furthermore, nonspiking LNs with weak active membrane properties that do not generate Na^+^ driven action potentials have been described in both holo- and hemimetabolous insect species (Husch et al., 2009a, 2009b; Tabuchi et al., 2015). While their functional role for odor information processing is not clear yet, it is plausible to assume that they are functionally highly relevant since they have been found across different insect species. In *P. americana*, the nonspiking LNs are named type II LNs (Fusca et al., 2013; Husch et al., 2009a, 2009b). They exist in two main types (type IIa and type IIb LNs) with different active membrane properties. Type IIa LNs have strong Ca^2+^ mediated active properties and respond to odorant stimulation with patterns of excitation and inhibition. A subset of type IIa LNs is cholinergic and can generate Ca^2+^ driven spikelets (type IIa1). Type IIb LNs respond with slow, sustained depolarizations.

Based on their functional and morphological properties, it can be hypothesized that they are mostly involved in intraglomerular signaling since the graded changes in membrane potentials can only spread within the same or electrotonically close glomeruli, as was proposed for nonspiking LNs in the rabbit olfactory bulb (Bufler et al., 1992a).

This study’s rationale was based on the previously reported structural and functional differences between distinct LN types in the cockroach AL (Fusca et al., 2013, 2015; Husch et al., 2009a, 2009b; Pippow et al., 2009). Spiking type I LNs are GABAergic, inhibitory, and innervate many but not all glomeruli. While some glomeruli are densely innervated, others are more sparsely or not at all innervated (Figure 1A). It has been considered that this reflects an organization with distinctive input and output glomeruli (Galizia and Kimmerle, 2004; Husch et al., 2009a; Wilson and Laurent, 2005). In this model, synaptic input is integrated and triggers action potential firing. The action potentials propagate to the innervated glomeruli and provide a defined glomeruli array with inhibitory synaptic input. Glomeruli can interact independently of their spatial and electrotonic distance. In this scenario, one would expect that odor-evoked glomerular Ca^2+^ signals are dominated by Ca^2+^ influx through voltage-gated channels that are activated by the action potentials. Thus, odor induced Ca^2+^ signals should be detectable and comparable in all innervated glomeruli.

**Figure 1.**
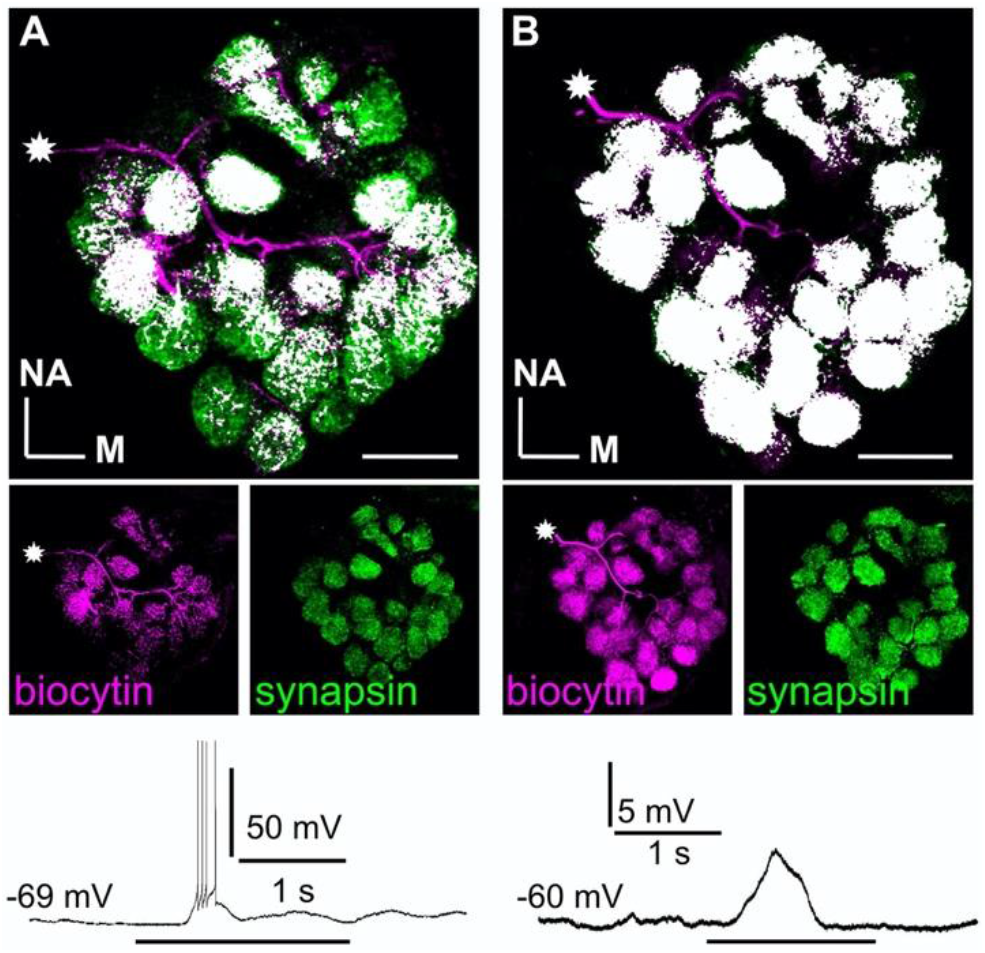
Branching patterns and odorant responses of spiking and nonspiking local interneurons. A spiking type I (**A**) and a nonspiking type II local interneuron (**B**) that were labeled with biocytin/streptavidin via the patch pipette. The glomeruli were visualized by synapsin-LIR. (**A**) Type I local interneuron. 13 µm stack of optical sections. The neuron innervates many but not all glomeruli and generates action potentials to an odorant stimulus (benzaldehyde). (**B**) Type II local interneuron. 15 µm stack of optical sections. The neuron innervated all glomeruli and responded to the odorant (benzaldehyde) with a graded depolarization. The stars mark the locations of the somata. Biocytin/streptavidin, magenta; synapsin-LIR, green; double-labeled pixels, white. NA: anterior, M: medial. Scale bars = 100 µm.

In contrast, nonspiking type II LNs have very similar branching patterns in all glomeruli (Figure 1B, Husch et al., 2009b), suggesting that both input and output can occur in every glomerulus. Due to the receptor and sensillum type-specific input configuration of the glomeruli in the AL (Fujimura et al., 1991; Watanabe et al., 2012), synaptic input during olfactory stimulation typically occurs only in a limited number of glomeruli (Sachse et al., 1999; Silbering et al., 2011). The resulting stimulus-evoked graded postsynaptic potentials can only spread within the same glomerulus or electrotonically nearby glomeruli. Since these neurons cannot generate Na^+^-driven action potentials, we hypothesize that the Ca^2+^ signals are dominated by odorant evoked Ca^2+^ influx through excitatory ligand-gated channels (Oliveira et al., 2010).

This study investigated the role of spiking and nonspiking LNs for inter- and intraglomerular signaling during olfactory information processing. To this end, we combined whole-cell patch clamp recordings with Ca^2+^ imaging to analyze the local Ca^2+^ dynamics of neurites in individual glomeruli as an indicator of signal processing in single LNs. The recordings were performed in the AL of the cockroach *Periplaneta americana*. This is an experimental system in which the olfactory system’s circuitry has been analyzed in great detail on the physiological (Bradler C, 2016; Ernst and Boeckh, 1983; Husch et al., 2009a, 2009b; Lemon and Getz, 1997, 1998, 2000; Nishino et al., 2012, 2018; Paeger et al., 2017; Paoli et al., 2020; Pippow et al., 2009; Strausfeld and Li, 1999; Warren and Kloppenburg, 2014; Watanabe et al., 2017), biochemical (Distler, 1989, 1990; Fusca et al., 2013, 2015; Neupert et al., 2012, 2018), and structural/ ultrastructural levels (Distler and Boeckh, 1997a, 1997b; Distler et al., 1998; Malun, 1991a, 1991b; Malun et al., 1993; Nishino et al., 2015; Watanabe et al., 2010), thus contributing very successfully to understanding olfactory information processing principles.

## RESULTS

Local interneurons of the insect AL are a heterogeneous group of neurons, consisting of different neuronal subpopulations with clearly defined, sometimes fundamentally different functional phenotypes. To study the role of spiking type I LNs and nonspiking type II LNs for inter and intraglomerular signaling, we combined whole-cell patch clamp recordings, Ca^2+^ imaging, and single cell labeling. This way, the cells were unequivocally identified by their physiological and morphological characteristics. In the investigated LNs, we measured the intracellular Ca^2+^ dynamics of neurites during olfactory stimulation simultaneously in many individual glomeruli. To determine differences in the odor-induced Ca^2+^ signals between individual glomeruli, tuning curves were constructed from the odor-evoked glomerular Ca^2+^ signals by normalizing them to the maximum signal amplitude of each glomerulus. Overall, this study is based on 17 recordings of type I LNs and 18 recordings of type II LNs. In each recorded LN, between 10 and 25 distinct glomeruli could be identified and individually imaged and analyzed.

In the first set of experiments, we showed that the odor-induced Ca^2+^-dynamics were highly reproducible when the antennae were repeatedly stimulated with the same odorant (Figure 2). This is in line with previous electrophysiological studies, in which LNs responded very reproducibly to repeated olfactory stimulations (Husch et al., 2009a, 2009b; Olsen and Wilson, 2008). Hence, in subsequent experiments, we analyzed single-sweep optophysiological recordings rather than averaged data.

**Figure 2.**
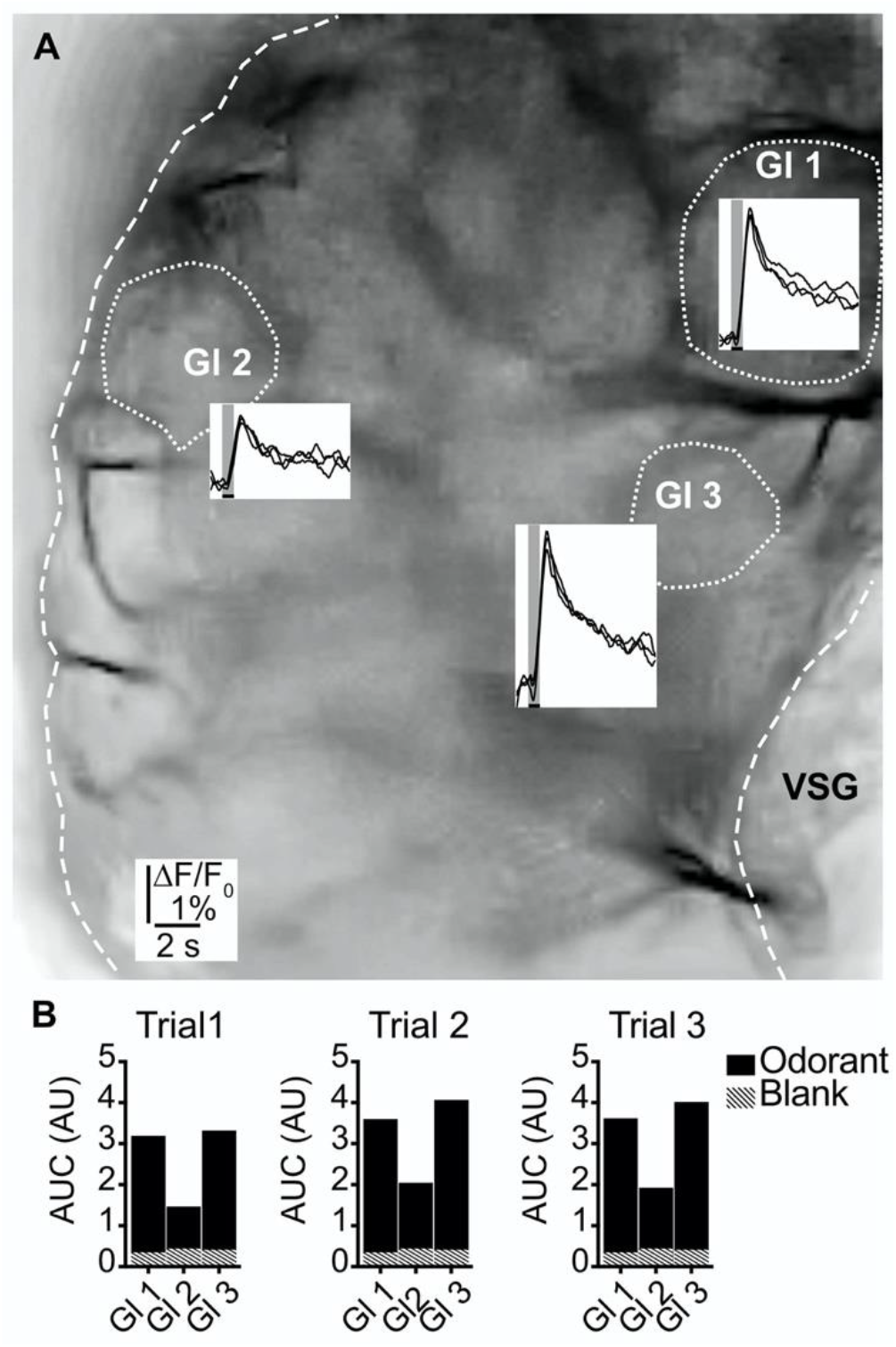
Odorant induced glomerular calcium signals are reproducible. (**A**) Transmitted light image of an investigated antennal lobe. The dotted lines mark the recorded glomeruli, and the insets show overlays of Ca^2+^ responses from three trials with the same odorant (hexanol). The grey bars mark the 500 ms odorant stimuli. (**B**) Areas under the curves of the Ca^2+^ signals that are shown in (A). The first three seconds after stimulus onset were analyzed. Hatched bars represent control signals to blank stimuli. AUC: area under the curve, Gl: glomerulus, VSG: ventrolateral somata group.

### Uniform glomerular odor responses in spiking type I LNs

All recorded type I LNs displayed characteristic morphological features, i.e., arborizations in multiple glomeruli with varying neurite densities between glomeruli (Figure 3A). Electrophysiologically, type I LNs reacted to odor stimulation of the antennae with odorant specific patterns of overshooting action potentials (Figure 3B). The glomerular Ca^2+^ signals were time-locked with the electrophysiological responses. While the absolute amplitudes of the Ca^2+^ signals during a given odorant varied between individual glomeruli, the time course and overall structure of the Ca^2+^ signals were very similar in all recorded glomeruli for a particular odorant (Figure 3B, Figure 3-figure supplement 1), resulting in identical tuning curves for all glomeruli of a given neuron (Figure 3C). Accordingly, tuning curves of all imaged glomeruli of a given neuron always correlated with coefficients of ∼ 1, with a mean correlation coefficient across all investigated spiking LNs of r=0.96 ± 0.03 (N=17, Figure 3D-F).

**Figure 3.**
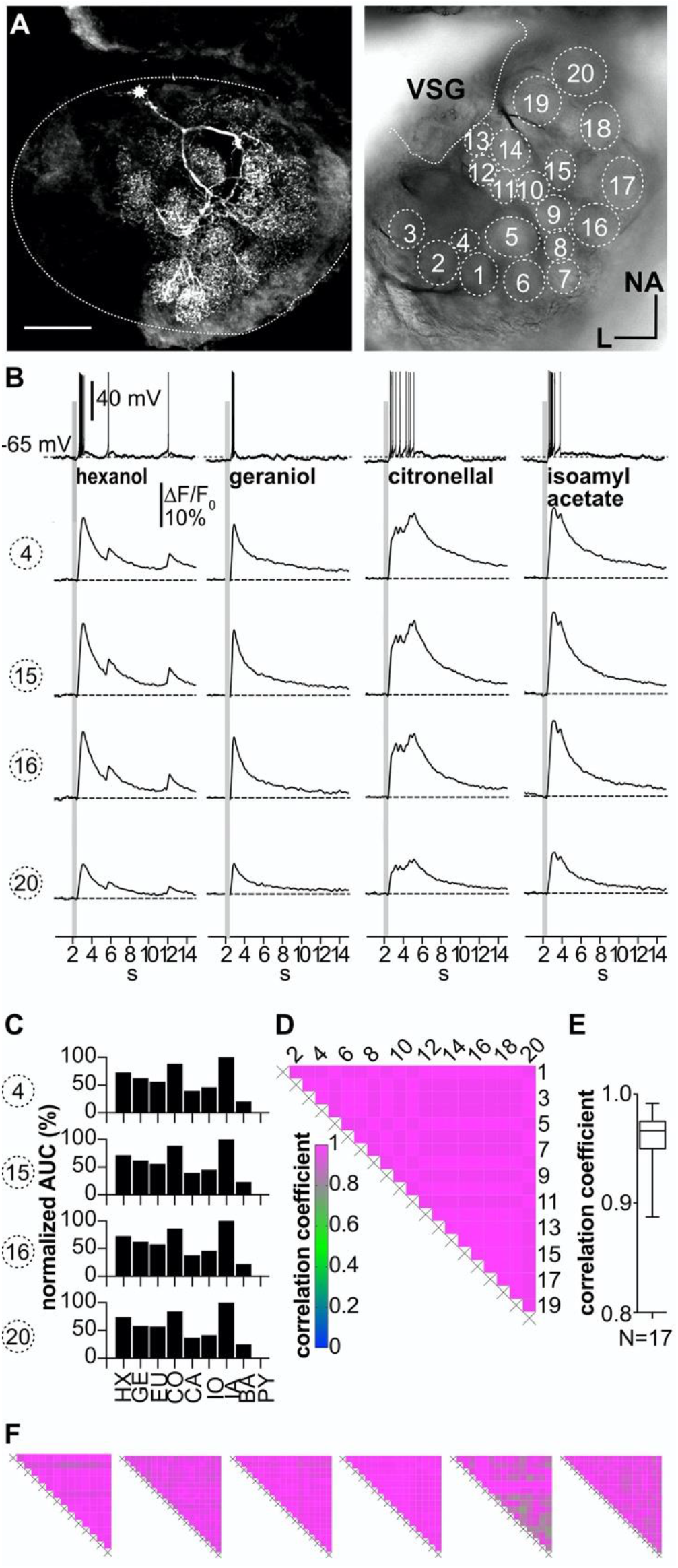
Ca^2+^ imaging in type I LNs shows uniform glomerular odor responses. (**A**) Left: Biocytin/streptavidin labeled type I LN. The AL is outlined by the dotted line. The star marks the position of the soma. Scale bar: 100 µm. Right: Transmitted light image of the same AL while the neuron was recorded. Orientation applies to both images. The outlined glomeruli mark the regions of interest (individual glomeruli) that were individually analyzed. (**B**) Electrophysiological responses to four odorants (top traces) and the corresponding Ca^2+^ dynamics of four glomeruli that are marked in (A). Gray bars represent the 500 ms odorant stimuli. The neuron responded to different odorants with odorant specific spike trains. The time courses of the Ca^2+^ signals were similar in all glomeruli for a given odorant. (**C**) Tuning curves of glomerular responses. Areas under the curves of the odorant evoked glomerular Ca^2+^ signals (first 3 seconds after stimulus onset) were calculated for a set of nine odorants and normalized to the maximum response in the respective glomerulus. Every glomerulus responded most strongly to isoamyl acetate and least to benzaldehyde. (**D**) Heatmap showing the correlations between the glomerular tuning curves of every imaged glomerulus. Numbers correspond to the glomeruli in (A). All tuning curves were well correlated with coefficients of ∼1 (nonparametric Spearman correlation). (**E**) Mean correlation coefficient across all investigated type I LNs was 0.96 ± 0.03 (N = 17). (**F**) Heatmaps of correlations between glomerular tuning curves from six additional type I LNs. HX: hexanol, GE: geraniol, EU: eugenol, CO: citronellal, CA: citral, IO: ionone, IA: isoamylacetate, BA: benzaldehyde, PY: pyrrolidine.

### Suppression of action potential firing in type I LNs decreases correlations of glomerular Ca^2+^ signals

We hypothesized that the observed Ca^2+^ signals in type I LNs mainly reflect the voltage-dependent Ca^2+^ influx induced by propagated action potentials. To test this hypothesis, we used two approaches to prevent the neurons from spiking. The neurons were hyperpolarized to membrane potentials between −80 mV to −100 mV (Figure 4A, top trace) or firing was suppressed by intracellularly blocking Na^+^ channels with QX-314. When the generation of action potentials is inhibited, the remaining Ca^2+^ signals should mainly reflect Ca^2+^ influx via ligand-gated channels (e.g., cholinergic receptors, Oliveira et al., 2010).

**Figure 4.**
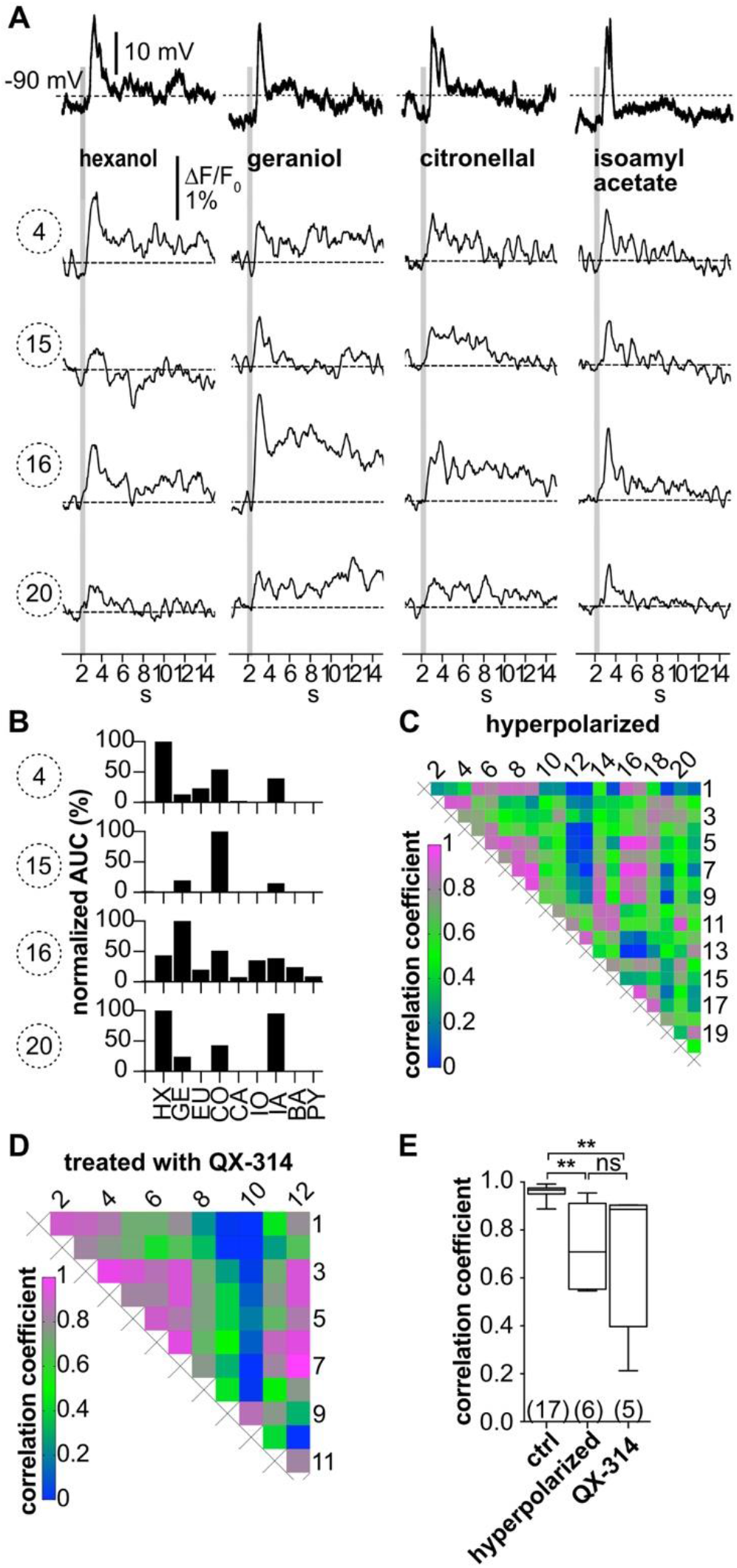
Hyperpolarization below the action potential threshold and pharmacological block of action potential firing prevent the correlation between odorant induced glomerular Ca^2+^ signals. Data in (A) - (C) are taken from the same type I LN as in Figure 3. (**A**) Electrophysiological responses to odorants (top traces) and corresponding Ca^2+^ dynamics in the same four glomeruli as shown in Figure 3. The neuron was hyperpolarized to prevent the generation of action potentials upon stimulation with odorants. Electrophysiologically, the neuron responded with odorant specific graded depolarizations. The high correlation of the glomerular Ca^2+^ signals shown in Figure 3 was inhibited. (**B**) The tuning curves of the glomerular responses (for details see Figure 3C) varied considerably, whereby the odorant that triggered the maximum Ca^2+^ signal in each individual glomerulus was different for each glomerulus. (**C**) Heatmap demonstrating the heterogeneous correlations between glomerular tuning curves. Numbers correspond to glomeruli in Figure 3A. Correlation coefficients ranged between 0 and 0.95 (median = 0.56). (**D**) Heatmap demonstrating the variable correlations between glomerular tuning curves of a neuron that was treated with the intracellular Na_V_ channel blocker QX-314. Correlation coefficients ranged between 0 and 1 (median = 0.69). (**E**) Mean correlation coefficients of hyperpolarized (0.73 ± 0.19, N = 6, p = 0.0024) and QX-314 treated (0.71 ± 0.28, N = 5, p = 0.0067) type I LNs were significantly decreased compared to the control group (Kruskal-Wallis and Dunn’s multiple comparisons test). Hyperpolarized and QX-314 treated type I LNs were not significantly different (p > 0.9999). Abbreviations as in Figure 3F.

When action potential firing was prevented by hyperpolarization, odor stimulation still elicited Ca^2+^ signals in the glomeruli. Besides a reduction in amplitude, the uniformity of the Ca^2+^ signals between different glomeruli disappeared. In turn, the tuning curves of the individual glomeruli became different from each other (Figure 4B and Figure 4–figure supplement 1). This is quantitatively reflected in the glomerulus-specific odorant responses and the diverse correlations between the glomerular tuning curves, resulting in a decreased mean correlation coefficient across all hyperpolarized type I LNs of r=0.73 ± 0.19 (N=6, Figure 4C-E). Similar results were obtained when AP firing was suppressed by intracellularly blocking Na^+^ channels with QX-314 (r=0.71 ± 0.28, N=5, Figure 4D,E). Differences in mean correlation coefficients were significant between control and hyperpolarized type I LNs (p=0.002) as well as between control and type I LNs that were treated with QX-314 (p=0.007). Mean correlation coefficients of hyperpolarized and QX-314 treated type I LNs were not significantly different (p>0.999).

Taken together, our results are in line with the conception that type I LNs integrate and transform their synaptic input to action potential firing to provide inhibitory synaptic input to neurons in a defined array of glomeruli. Our results also provide physiological evidence that an individual type I LN receives excitatory input not only in one but in several glomeruli, which is in line with previous structural- and ultrastructural studies that reported evidence for both pre- and postsynaptic profiles in individual glomeruli (Berck et al., 2016; Distler and Boeckh, 1997b; Mohamed et al., 2019).

### Variable glomerular odor responses in nonspiking type II LN

In contrast to the uniform Ca^2+^ dynamics during odor stimulation in type I LNs, we observed highly heterogeneous Ca^2+^ dynamics between the individual glomeruli in most type II LNs (Figure 5A-J). All recorded type II LNs had the cell type-specific morphology characterized by innervation of all glomeruli with similar neurite densities in all glomeruli of a given neuron (Figure 5A,E). All type II LNs typically reacted to the olfactory stimulation with graded changes in membrane potential. (Figure 5B,F, top traces). The amplitudes of the corresponding Ca^2+^ signals were in the range of the signals of type I LNs after the suppression of their action potentials. While the electrophysiological responses to different odorants were similar in a given neuron, the corresponding glomerular Ca^2+^ signals were odor specific and varied between glomeruli, resulting in distinct, glomerulus specific tuning curves (Figure 5C-H and Figure 5–Figure supplement 1), which was also directly evident in a rather low degree of correlation (Figure 5I; r=0.53 ± 0.23, N=18). However, the correlation between tuning curves of individual glomeruli in a given neuron differed among type II LNs. While in most nonspiking LNs, the majority of glomeruli was individually tuned, in 8 out of 18 neurons, groups of similarly tuned glomeruli were found. This is shown in the heatmaps showing highly correlated Ca^2+^ signals in groups of glomeruli as well as glomeruli that were not correlated (Figure 5G,H,J and Figure 5–figure supplement 2). Mechanistically, this could be caused by similar input to several glomeruli (Watanabe et al., 2012) or by coordinated activity, e.g., via spikelets that were observed in a sub-type of nonspiking neurons (type IIa1 LNs, Fusca et al., 2013).

**Figure 5.**
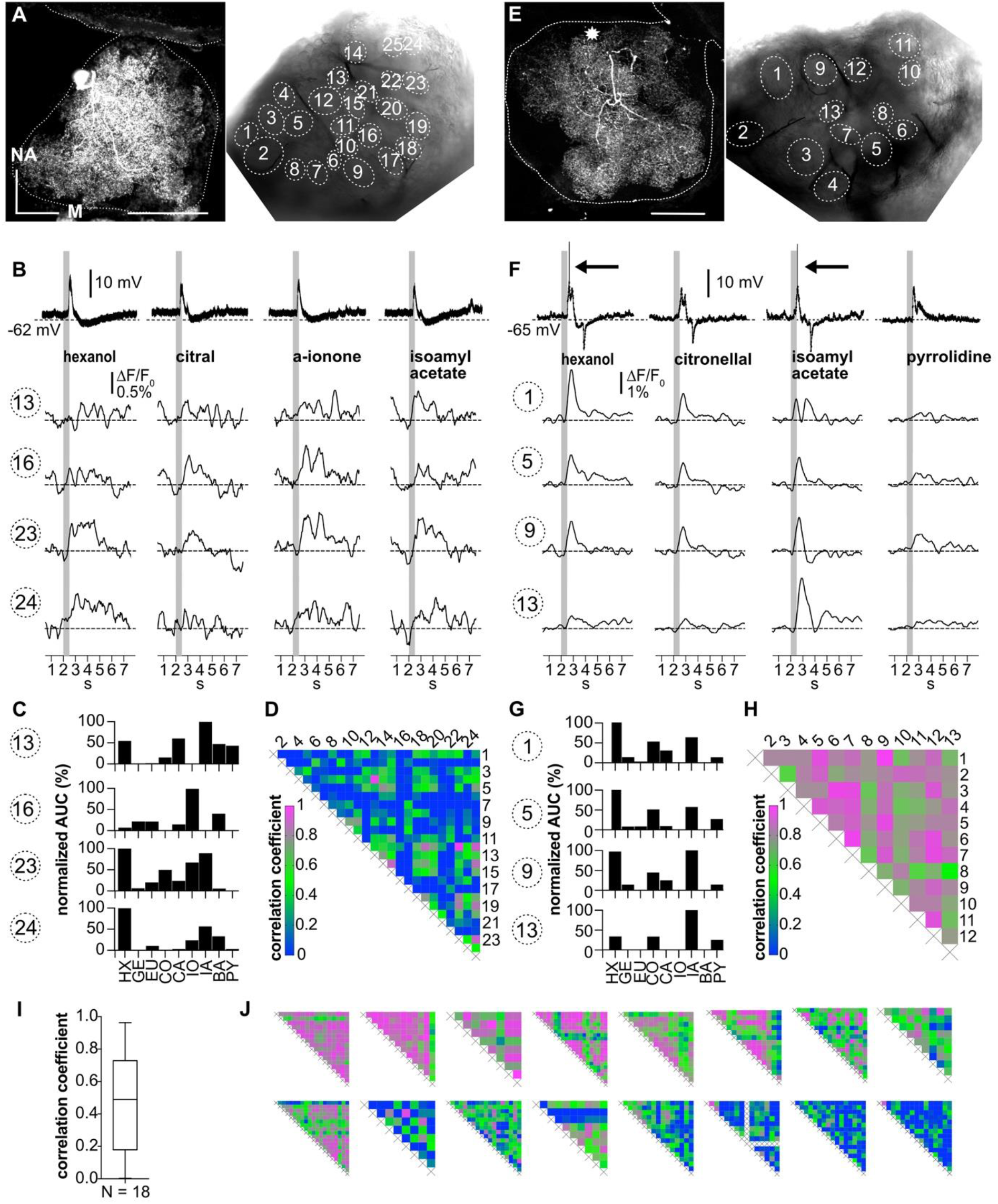
Ca^2+^ imaging of type II LNs shows heterogeneous glomerular odorant responses. Data from a type IIb (**A**-**D**), and a type IIa LN (**E**-**H**). (**A, E**) Left: Biocytin/streptavidin stainings of the investigated type II LNs. The ALs are outlined by the dotted lines. The position of the soma in (E) is marked by the star. Scale bar: 100 µm. Right: Transmitted light images of the same ALs during the experiment. Outlined glomeruli were marked as regions of interest and individually analyzed. The orientations of the left and right images are similar. (**B, F**) Electrophysiological responses to four odorants (top traces) with the corresponding Ca^2+^ dynamics of four glomeruli that are marked in the images shown in A and E. Gray bars represent the 500 ms odorant stimuli. (**B-D**) Type IIb LN. (**B**) The neuron responded similarly to the different odorants with graded depolarizations that were followed by slow hyperpolarizations. The time course and amplitude of the corresponding Ca^2+^ signals varied in different glomeruli for the different odorants. (**C**) Tuning curves of glomerular Ca^2+^ signals (for details, see Figure 3C). The tuning curves of the different glomeruli varied considerably, while the maximum response was induced by different odorants in the different glomeruli. Some glomeruli were narrowly tuned (e.g., glomerulus 16); others were broadly tuned (e.g., glomerulus 23). (**D**) Heatmap showing the correlations between glomerular tuning curves of every imaged glomerulus. Numbers correspond to glomeruli shown in (A). Correlations between glomerular tuning curves were mostly low, with coefficients ranging between 0 and 0.96 (median = 0.15). (**F**-**H**) Type IIa LN. (**F**) The neuron responded similarly to different odorants with graded depolarizations that could include spikelets (e.g. hexanol, isoamylacetate, arrows mark the spikelets), whereas the time course and amplitude of the corresponding Ca^2+^ signals mostly varied between different glomeruli for different odorants. (**G**) Tuning curves of the glomerular Ca^2+^ signals shown in (F) (for details, see Figure 3C). Groups of glomeruli showed similar tuning curves (e.g., glomeruli 1, 5, and 9), while other glomeruli were individually tuned (e.g., glomerulus 13). (**H**) Heatmap showing correlations between glomerular tuning curves of every imaged glomerulus. Numbers correspond to the glomeruli marked in (E). Glomerular tuning curves correlated strongly in a subset of glomeruli, while the correlation was low between other glomeruli. Coefficients ranged between 0.45 and 0.96 (median = 0.76). (**I**) Mean correlation coefficients across all investigated type II LNs were 0.53 ± 0.23 (N = 18). (**J**) Heatmaps of correlations between glomerular tuning curves from all additional type II LNs in descending order of mean correlation coefficient. Abbreviations as in Figure 3F.

## DISCUSSION

Processing of sensory input by networks of spiking and nonspiking interneurons is a common principle in both invertebrate and vertebrate sensory systems, e.g., structuring the signal pathway from sensory neurons (tactile hairs) to intersegmental and motor neurons in the insect thoracic ganglion (Burrows, 1989; Pearson and Fourtner, 1975) and the mammalian olfactory bulb (Bufler et al., 1992b, 1992a; Wellis and Scott, 1990) and retina (Diamond, 2017). Nevertheless, in many systems, the role of nonspiking neurons is not well understood.

Local interneurons are key components of the insect olfactory system. They have fundamentally different functional phenotypes suggesting different tasks during odor information processing. To help elucidate mechanisms of odor processing on the level of individual LNs, this study assessed local Ca^2+^ dynamics in distinct functional compartments (ramifications in individual glomeruli) of spiking type I and nonspiking type II LNs during olfactory information processing. To this end, individual LNs were analyzed by combined whole-cell patch clamp recordings and Ca^2+^ imaging. Local Ca^2+^ dynamics are likely to reflect the role of LNs in odor information processing, i.e., for their potential role in intra- and interglomerular signaling, which depends crucially on signal propagation throughout individual LNs. In line with the electrophysiological properties, we found odorant evoked Ca^2+^ signals that were homogeneous across the whole cell in spiking type I LNs and odor and glomeruli specific Ca^2+^ signals in nonspiking type II LNs. This is reflected in the highly correlated tuning curves in type I LNs and low correlations between tuning curves in type II LNs. In the following, we discuss whether and how this is consistent with previous studies suggesting that spiking type I LNs play a role in lateral, interglomerular signaling and why this study assigns a role to nonspiking LNs in local, intraglomerular signaling.

### Interglomerular signaling via spiking type I LNs

Processing of olfactory information in the AL involves complex interactions between the glomerular pathways and between different AL neurons. Previous studies in different insect species have suggested that GABAergic LNs can mediate lateral inhibition by providing inhibitory synaptic input to defined odor specific arrays of glomeruli. This hypothesis is in agreement with the current study, where spiking type I LNs showed odorant specific glomerular Ca^2+^ dynamics, which were always uniform in every imaged glomerulus. As the glomerular signals in these neurons correlated very well with the electrophysiological activity, it is plausible that synaptic inputs to one or a few glomeruli are integrated and result in the firing of action potentials, which propagate to the neurites of all innervated glomeruli where they induce highly correlated voltage-activated Ca^2+^ signals. Since type I LNs express GABA-like immunoreactivity and provide inhibitory input to uPNs and other LNs in all innervated glomeruli (Distler, 1989; Distler and Boeckh, 1997c; Husch et al., 2009a; Warren and Kloppenburg, 2014), these neurons are likely part of an inhibitory network that mediates lateral inhibition and contrast enhancement (Sachse and Galizia, 2002; Wilson and Laurent, 2005).

However, it is important to consider that the experiments with suppressed AP firing showed not well correlated odor-induced Ca^2+^ signals, i.e., tuning curves in the individual glomeruli of type I LNs. This is in line with the hypothesis that odor induced Ca^2+^ signals under control condition originate mostly from action potential induced Ca^2+^ influx via voltage-activated Ca^2+^ channels. These data also show that the hypothesized polar organization of type I LNs with strictly defined (and separated) input and output glomeruli is not entirely correct. When type I LNs were prevented from spiking by hyperpolarization or intracellular block of Na^+^ channels, we observed distinct, glomeruli specific Ca^2+^ dynamics during odor stimulation. It is likely that these signals originate from Ca^2+^ influx through Ca^2+^ permeable excitatory receptors such as cholinergic receptors, suggesting that each neuron can potentially receive excitatory olfactory input in any innervated glomerulus. This notion agrees with previous studies in the fly, which suggested that individual GABAergic LNs receive broad, but not uniform, spatial patterns of excitation by either OSNs or PNs (Wilson and Laurent, 2005).

### Inter- and intraglomerular signaling via nonspiking type II LN

The nonspiking LNs that have been described in the AL of insects typically innervate all glomeruli (Husch et al., 2009a, 2009b; Tabuchi et al., 2015). In most of these neurons, we observed highly heterogeneous Ca^2+^ dynamics between the individual glomeruli resulting in distinct tuning curves for the individual glomeruli. These heterogeneous and glomerulus specific Ca^2+^ dynamics imply that type II LNs have distinct functional domains that are (more or less) independent from each other. Accordingly, the majority of nonspiking type II LNs might contribute to microcircuits within glomeruli and mediate intraglomerular signaling rather than interconnecting multiple glomeruli.

Intraglomerular circuits are known from the mouse or rat olfactory bulb (for review see Ennis et al., 2015), where periglomerular, external tufted, and short axon cells interact to modulate the output of Mitral/ Tufted cells (Aungst et al., 2003; Liu et al., 2016; Najac et al., 2015; Wachowiak and Shipley, 2006). Periglomerular cells are uniglomerular local interneurons that mediate intraglomerular synaptic signaling. Unlike periglomerular cells, cockroach type II LNs innervate all glomeruli. Still, as these LNs have only weak active membrane properties, postsynaptic potentials just spread within the same glomerulus. Therefore, these neurons could serve similar purposes, and few omniglomerular type II LNs could perform similar functions as many PG uniglomerular cells.

In addition, it is important to consider that nonspiking type II LNs are not a homogenous neuron population (Fusca et al., 2013; Husch et al., 2009b). In a subpopulation of type II LNs, we observed correlated Ca^2+^ dynamics in subsets of glomeruli. These neurons typically responded to odorant stimulations with strong depolarizations, including spikelets, which apparently can propagate, at least to some extent, to a set of glomeruli. Since this subpopulation of nonspiking type II LNs (type IIa1 LNs) was previously shown to be cholinergic (Fusca et al., 2013; Neupert et al., 2018), they are likely excitatory. The intrinsic electrophysiological properties of the cholinergic type IIa LNs suggest that they might be part of an excitatory network, which activates neurons in specific sets of glomeruli. This hypothesis is in line with previous studies in the fruit fly, where excitatory LNs, while being multiglomerular, only activate specific glomeruli, thereby providing distinct arrays of glomeruli with excitatory input and distributing odor-evoked activity over an ensemble of PNs (Das et al., 2017; Huang et al., 2010; Olsen et al., 2007; Root et al., 2007; Shang et al., 2007; reviewed in Wilson, 2013).

While type IIa1 are cholinergic and type II LNs generally express multiple neuropeptides (Fusca et al., 2015; Neupert et al., 2012, 2018), the primary transmitter of most type II LNs is yet to be revealed. One candidate is glutamate, which is an inhibitory transmitter in the *Drosophila* AL (Liu and Wilson, 2013) and in cockroach metathoracic motor neurons (Sattelle, 1992).

We conclude that in the cockroach AL, sensory inputs are processed and computed in inter- and intraglomerular circuits which are formed by spiking type I and nonspiking type II LN.

## METHODS

### Animals and materials

*P. americana* were reared in crowded colonies at 27 °C under a 13 : 11 h light/ dark photoperiod regimen, on a diet of dry rodent food, oatmeal, and water. The experiments were performed with adult males. Unless stated otherwise, all chemicals were obtained from Applichem (Darmstadt, Germany) or Sigma-Aldrich (Taufkirchen, Germany) and had the purity level ‘pro analysis’.

### Intact brain preparation

The brain preparation leaving the entire olfactory network intact has been described previously (Demmer and Kloppenburg, 2009; Husch et al., 2009a; Kloppenburg et al., 1999). Animals were anesthetized by CO_2_, placed in a custom-built holder, and the head was immobilized with tape (tesa ExtraPower Gewebeband, Tesa, Hamburg, Germany). The head capsule was opened by cutting a window between the two compound eyes and the antennae’s bases. The brain with its antennal nerves and attached antennae was dissected in extracellular saline (see below) and pinned in a Sylgard-coated (Dow Corning Corp., Midland, Michigan, USA) recording chamber. To get access to the recording site, we desheathed parts of the AL using fine forceps, and preparations were enzymatically treated with a combination of papain (0.3 mg·ml^-1^, P4762, Sigma) and L-cysteine (1 mg·ml^-1^, 30090, Fluka) dissolved in extracellular saline (∼3 min, room temperature (RT), ∼24°C). For electrophysiological recordings, the somata of the AL neurons were visualized with a fixed stage upright microscope (AxioExaminer, Carl Zeiss, Jena, Germany) using a 20x water-immersion objective (20x W Apochromat, NA=1) with a 4x magnification changer, and infrared differential interference contrast optics (Dodt and Zieglgansberger, 1994).

### Identification of antennal lobe neurons

The prerequisite to study the physiology of identified neurons is the unequivocal identification of neuron types. The identification was performed as described by Fusca et al., 2013. Briefly, AL neurons were first pre-identified by the size and location of their somata. Recordings were performed under visual control from cell bodies in the ventrolateral somata group (VSG, Distler, 1989), where different neuron types are located in separated clusters. This pre-identification has a high success rate for the major neuron types (>90%), and was verified in each case by the physiological and morphological characterization during and after the recording using the following criteria: Two main LN types were identified by their distinctive physiological properties: 1) spiking type I LNs that generated Na^+^ driven action potentials upon odor stimulation and, 2) nonspiking type II LNs, in which odor stimulation evoked depolarizations, but no Na^+^ driven action potentials (Husch et al., 2009a, 2009b). Type I LNs had arborizations in many, but not all glomeruli. The density of processes varied between glomeruli of a given type I LNs. Type II LN had processes in all glomeruli. The density and distribution of arborizations were similar in all glomeruli of a given type II LN, but varied between different type II LN. Two sub-types (type IIa and type IIb) can be distinguished by the branch patterns within the glomeruli, the size and branch pattern of low order neurites, odor responses, and active membrane properties (Husch et al., 2009b). Type IIa LNs had strong Ca^2+^ dependent active membrane properties and responded with odor specific elaborate patterns of excitation and periods of inhibition. In a subset of the type IIa neurons, which are cholinergic (type IIa1, Fusca et al., 2013) the excitation included Ca^2+^ driven ‘spikelets’ riding on the depolarization. In contrast, type IIb LNs responded mostly with sustained, relatively smooth depolarizations.

### Whole cell recordings

Whole-cell recordings were performed at RT following the methods described by Hamill et al., 1981. Electrodes with tip resistances between 2-3 MΩ were fashioned from borosilicate glass (inner diameter 0.86 mm, outer diameter 1.5 mm, GB150-8P, Science Products, Hofheim, Germany) with a vertical pipette puller (PP-830 or PC-10, Narishige, Japan). Recording pipettes were filled with intracellular solution containing (in mM): 218 K-aspartate, 10 NaCl, 2 MgCl_2_, 10 HEPES, and 0.8 Oregon Green 488 BAPTA-1 hexapotassium salt (OGB1, O6806, ThermoFisher Scientific, Waltham, MA, USA) adjusted to pH 7.2 with KOH. In some experiments, 2 mM lidocaine N-ethyl chloride (QX-314, #Q-150, Alomone, Jerusalem, Israel) was added to the intracellular solution. During the experiments, if not stated otherwise, the cells were superfused constantly with extracellular solution containing (in mM): 185 NaCl, 4 KCl, 6 CaCl_2_, 2 MgCl_2_, 10 HEPES, 35 D-glucose. The solution was adjusted to pH 7.2 with NaOH.

Whole-cell current clamp recordings were made with an EPC10 patch clamp amplifier (HEKA-Elektronik, Lambrecht, Germany) controlled by the program Patchmaster (version 2.53, HEKA-Elektronik) running under Windows. The electrophysiological data were sampled at 10 kHz. The recordings were low pass filtered at 2 kHz with a 4-pole Bessel-Filter. Compensation of the offset potential and capacitive currents was performed using the ‘automatic mode’ of the EPC10 amplifier. Whole-cell capacitance was determined by using the capacitance compensation (C-slow) of the amplifier. The liquid junction potential between intracellular and extracellular solution was also compensated (16.9 mV, calculated with Patcher’s-Power-Tools plug-in [https://www3.mpibpc.mpg.de/groups/neher/index.php?page=aboutppt] for Igor Pro 6 [Wavemetrics, Portland, Oregon]). Voltage errors due to series resistance (*R*_*S*_) were minimized using the RS-compensation of the EPC10. *R*_*S*_ was compensated between 60% and 70% with a time constant (τ) of 10 µs.

### Odor stimulation

To deliver the odorants, we used a continuous airflow system. Carbon-filtered, humidified air was guided across the antenna at a flow rate of ∼2 l·min^-1^ (‘main airstream’) through a glass tube (inner diameter 10 mm) that was placed perpendicular to and within 20-30 mm distance of the antennae. To apply odorants, 5 ml of odorant-containing solutions (either pure or diluted in mineral oil [M8410, Sigma]) were transferred into 100 ml glass vessels. Strips of filter paper in the odorant solution were used to facilitate evaporation. The concentration of each odorant was adjusted to match the vapor pressure of the odorant with the lowest value (eugenol). Dilutions were as follows: α-ionone 40.9% (I12409, Aldrich), +/− citral 24.2% (C83007, Aldrich), 1-hexanol 2.4% (52830, Fluka), benzaldehyde (418099, Aldrich) 2.2%, citronellal 8.7% (C2513, Aldrich), eugenol 100% (E51791, Aldrich), geraniol 73.7% (48799, Fluka), Isoamylacetate (112674, Aldrich) 13.7%, pyrrolidine 0.035% (83241, Fluka). The headspace of pure mineral oil was the control stimulus (blank). During a 500 ms stimulus, ∼17 ml of the vessel volume was injected into the main air stream. The solenoids were controlled by the D/A-interface of the EPC10 patch clamp amplifier and the Patchmaster software. Odorant-containing air was quickly removed from the experimental setup with a vacuum funnel (inner diameter 3.5 cm) placed 5 cm behind the antennae. To allow for sensory recovery, consecutive odorant stimulations of the same preparation were performed after intervals of at least 60 s with non-odorant containing air.

### Calcium Imaging

Odor evoked calcium dynamics were measured with the Ca^2+^ indicator OGB1 (see the intracellular solution), a single wavelength, high-affinity dye suitable to monitor fast intracellular Ca^2+^ signals. The imaging setup consisted of a Zeiss AxioCam/MRm CCD camera with a 1388×1040 chip and a Polychromator V (Till Photonics, Gräfelfing, Germany) that was coupled *via* an optical fiber into the Zeiss AxioExaminer upright microscope. The camera and polychromator were controlled by the software Zen pro, including the module ‘Physiology’ (2012 blue edition, Zeiss). After establishing the whole-cell configuration, neurons were held in current clamp mode, and a hyperpolarizing current (∼-200 pA) was injected for about 45-60 min to allow for dye loading. After loading, up to nine different odorants were applied as 500 ms pulses onto the ipsilateral antenna. Odor-induced Ca^2+^ transients in the OGB1-loaded neurons were monitored by images acquired at 488 nm excitation with 50 ms exposure time and a frame rate of ∼18 Hz. The emitted fluorescence was detected through a 500-550 nm bandpass filter (BP525/50), and data were acquired using 5×5 on-chip binning. Images were recorded in arbitrary units (AU) and analyzed as 16-bit grayscale images.

### Analysis of odor-evoked calcium signals

The analysis was performed offline using ImageJ (version 2.0.0-rc-64/1.51s) and Prism 7 (GraphPad, California, USA). Amplitudes and kinetics of the Ca^2+^ signals were calculated as means (in AU) of individual glomeruli, which were defined as the respective regions of interest (ROI). ROI were defined on transmitted light images of the investigated antennal lobes. The Ca^2+^ signals are given as relative fluorescence changes (*ΔF/F*_*0*_). To correct for bleaching, biexponential fits to the time courses of the glomerular Ca^2+^ signals during the blank stimulus, which lacked the odorant evoked Ca^2+^ influx, were used.

For statistical analysis of data obtained for the different cell types, nonparametric Kruskal-Wallis tests with Dunn’s multiple comparisons tests were performed in Prism 7. Correlation coefficients from matrices of glomerular tuning curves are given as nonparametric Spearman correlation r. A significance level of 0.05 was accepted for all tests. All calculated values are expressed as mean ± standard deviation.

### Single-cell and double-labeling and confocal microscopy

To label individual cells, 1% (w/v) biocytin (B4261, Sigma) was added to the pipette solution. After the electrophysiological recordings, the brains were fixed in Roti-Histofix (P0873, Carl Roth, Karlsruhe, Germany) overnight at 4°C. Subsequently, the brains were rinsed in 0.1 M phosphate buffered saline (PBS, 3 x 20 min and then for ∼12 h, RT). PBS contained (in mM) 72 Na_2_HPO_4_x2H_2_O, 28 NaH_2_PO_4_xH_2_O, resulting in pH 7.2. To facilitate streptavidin penetration, the samples were treated with a commercially available collagenase/dispase mixture (1 mg·ml^-1^, 269638, Roche Diagnostics, Mannheim, Germany) and hyaluronidase (1 mg·ml^-1^, H3506, Sigma-Aldrich) in PBS (1 h, 37°C), rinsed in PBS (3 x 10 min, 4°C) and then pre-incubated in blocking solution, consisting of PBS containing 1% (w/v) Triton X-100 (A1388, AppliChem) and 10% (v/v) normal goat serum (S-1000, Vector Labs, Burlingame, CA) for 1 h at RT. The brains were then incubated with *Alexa 633* conjugated streptavidin (1:400, S21375, Invitrogen, Eugene, OR) in PBS supplemented with 10% (v/v) normal goat serum for ∼12 h at 4°C, rinsed in PBS (3 x 10 min, RT), dehydrated, cleared, and mounted in methylsalicylate.

In some preparations, we used immunohistochemistry to label synapsin to mark the glomeruli. After pre-incubation in blocking solution and before the streptavidin incubation, these brains were incubated for 5 days at 4°C with a monoclonal primary mouse antibody against the presynaptic vesicle protein synapsin I (3C11, supernatant; obtained from the Developmental Studies Hybridoma Bank, University of Iowa, RRID: AB_528479), diluted 1:50 in blocking solution. Subsequently, the brains were rinsed in PBS-1% Triton X-100 (2×2h, RT), incubated in *Alexa 488* conjugated goat anti-mouse secondary antibody for 5 days at 4°C (1:200 in blocking solution, 115-545-062, Dianova, Hamburg, Germany) and rinsed in PBS-1% Triton X-100 (2×2h, RT) and PBS (3×10min, RT). 3C11 (anti-SYNORF1) was deposited to the DSHB by Buchner, E. (DSHB Hybridoma Product 3C11 (anti SYNORF1, Klagges et al., 1996)).

Fluorescence images were captured with confocal microscopes equipped with Plan-Apochromat 10x (numerical aperture 0.45) and Plan-Apochromat 20x (numerical aperture 0.75) objectives (LSM 510, Zeiss) or with a 20x objective (SP8, Leica Microsystems, Wetzlar, Germany) respectively. *Alexa 633* was excited at 633 nm, and emission was collected through a 650 nm long-pass filter. *Alexa 488* was excited at 488 nm, and emission was collected through a 505-530 nm bandpass filter. Confocal images were adjusted for contrast and brightness and overlaid in ImageJ. The final figures were prepared in Affinity Designer (Serif, Nottingham, UK).

## ROLE OF AUTHORS

Study concept and design: DF and PK. Acquisition of data: DF. Analysis and interpretation of data: DF and PK. Drafting the article: DF and PK.

## ACKNOWLEDGEMENTS

We thank Helmut Wratil for his excellent technical assistance. The work in PK’s laboratory was supported by grants from the Deutsche Forschungsgemeinschaft.

## FIGURES

**Figure 3 – Figure supplement 1.**
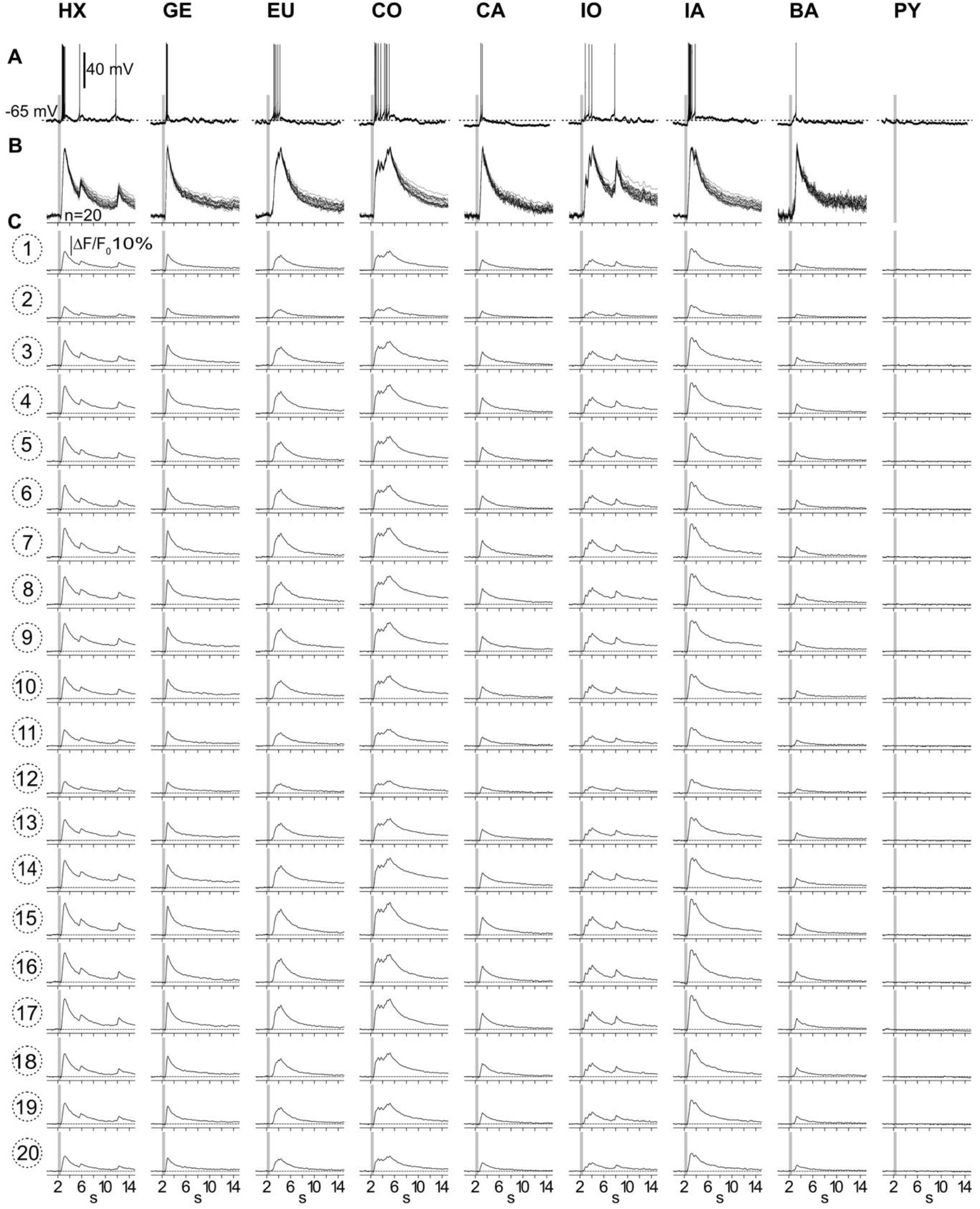
Ca^2+^ signals from all 20 imaged glomeruli. (**A**) Electrophysiological responses to all tested odorants. (**B**) Scaled overlays of the corresponding Ca^2+^ signals from all imaged glomeruli (n=20). The odorant induced Ca^2+^ signals were scaled to the same size. (**C**) Original odorant induced Ca^2+^ signals from all glomeruli.

**Figure 4 – Figure supplement 1.**
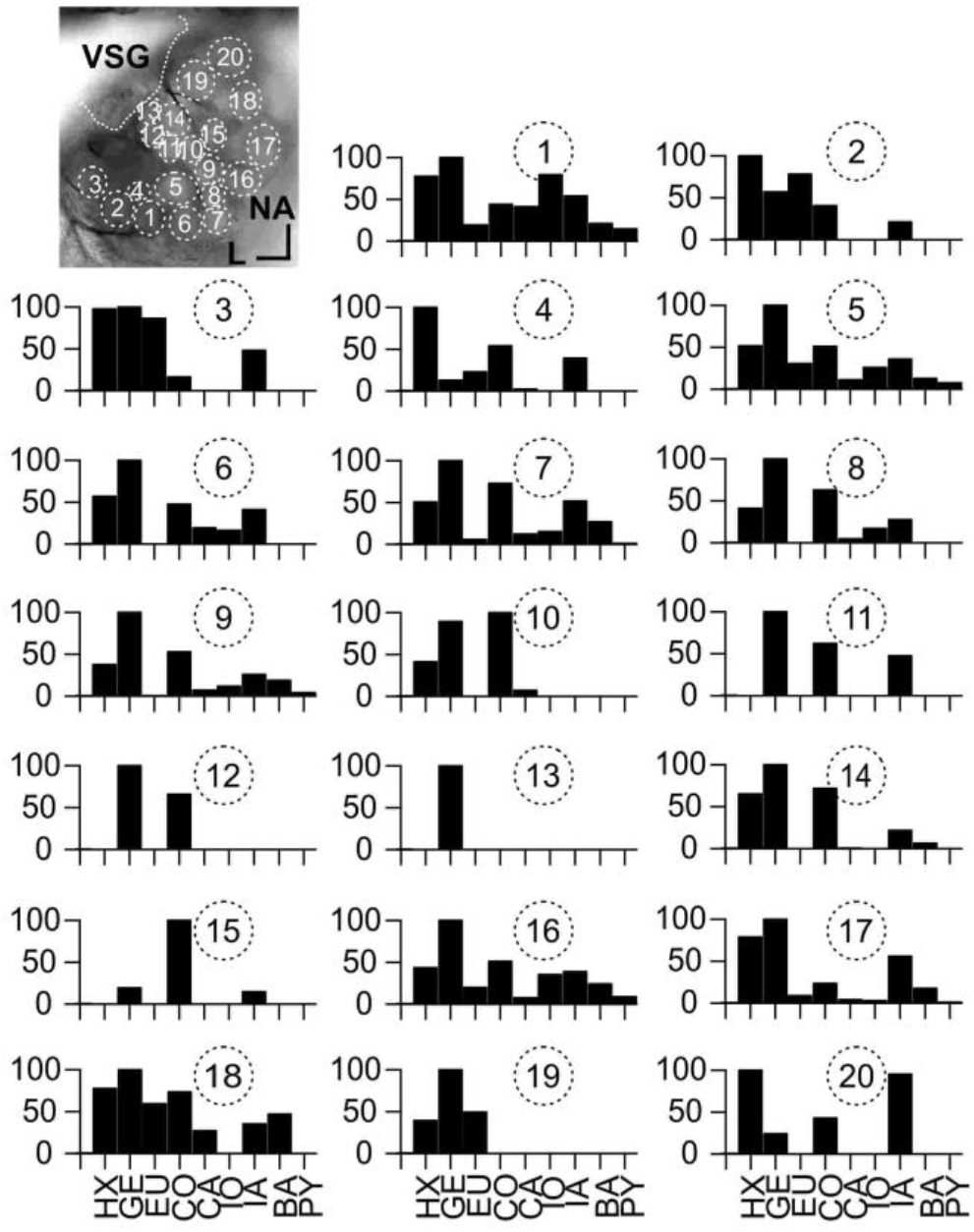
Tuning curves of all imaged glomeruli. Y-axes show normalized odor response. For details, see Figure 3C. Abbreviations as in Figure 3F.

**Figure 5 – Figure supplement 1.**
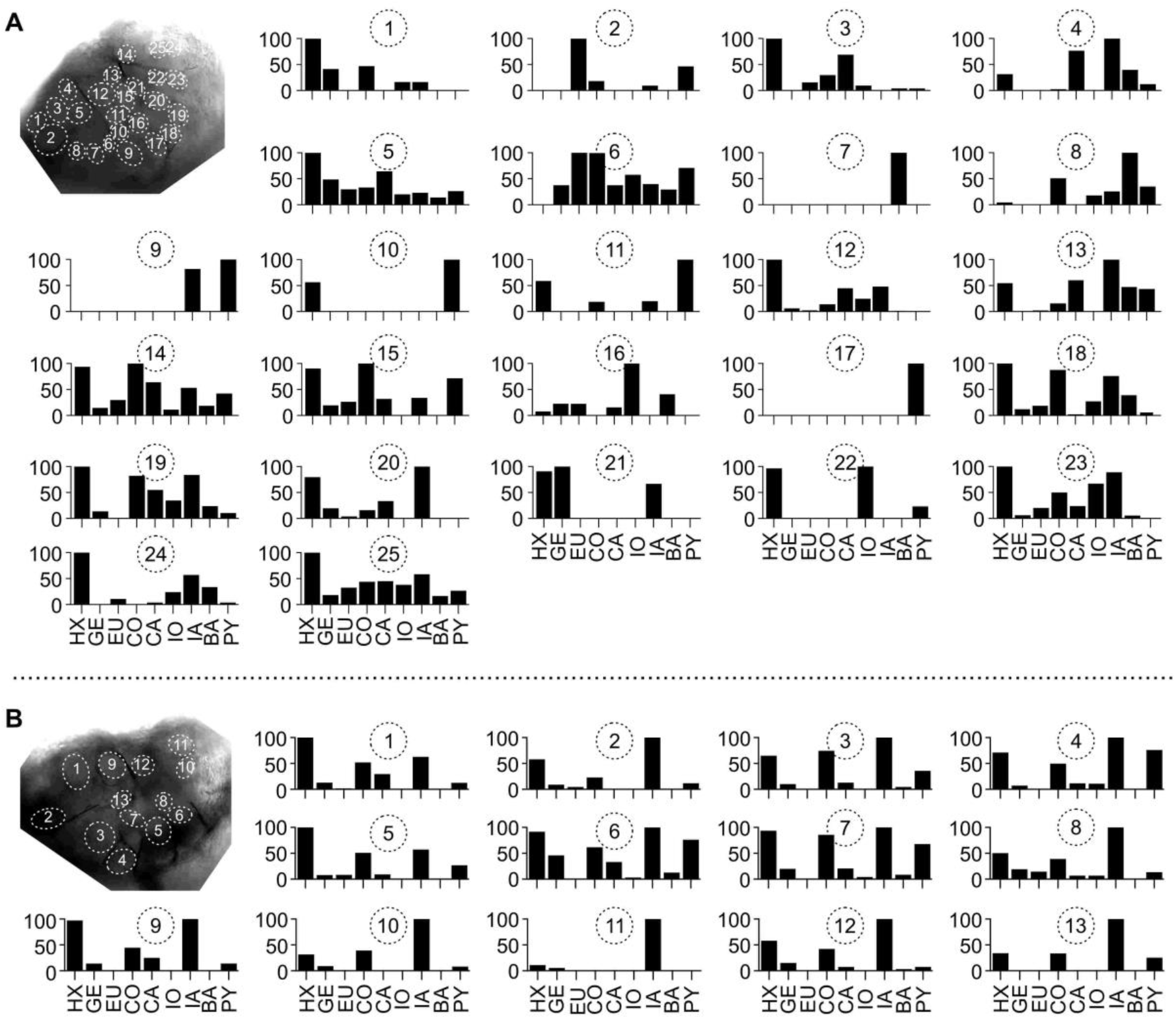
Glomerular tuning curves of all imaged glomeruli. Tuning curves of all glomeruli from the neurons shown in Figure 5A-D (**A**) and Figure 5E-H (**B**). Y-axes show normalized odor response. For details, see Figure 3C. Abbreviations as in Figure 3F.

## REFERENCES

Assisi, C., and Bazhenov, M. (2012). Synaptic inhibition controls transient oscillatory synchronization in a model of the insect olfactory system.Front. Neuroengineering 5, 7.

Assisi, C., Stopfer, M., and Bazhenov, M. (2012). Excitatory local interneurons enhance tuning of sensory information. PLoS Comput Biol 8, e1002563.

Aungst, J.L., Heyward, P.M., Puche, A.C., Karnup, S.V., Hayar, A., Szabo, G., and Shipley, M.T. (2003). Centre-surround inhibition among olfactory bulb glomeruli. Nature 426, 623–629.

Berck, M.E., Khandelwal, A., Claus, L., Hernandez-Nunez, L., Si, G., Tabone, C.J., Li, F., Truman, J.W., Fetter, R.D., Louis, M., et al. (2016). The wiring diagram of a glomerular olfactory system. ELife 5.

Berg, B.G., Schachtner, J., Utz, S., and Homberg, U. (2007). Distribution of neuropeptides in the primary olfactory center of the heliothine moth Heliothis virescens. Cell Tissue Res 327, 385–398.

Bradler C K.P., Warren B, Bardos V, Schleicher S, Klein A. (2016). Properties and physiological function of Ca2+-dependent K+ currents in uniglomerular olfactory projection neurons. J. Neurophysiol. 115, 2330–2340.

Bufler, J., Zufall, F., Franke, C., and Hatt, H. (1992a).Patch-clamp recordings of spiking and nonspiking interneurons from rabbit olfactory bulb slices: membrane properties and ionic currents. J. Comp. Physiol. [A] 170, 145–152.

Bufler, J., Zufall, F., Franke, C., and Hatt, H. (1992b). Patch-clamp recordings of spiking and nonspiking interneurons from rabbit olfactory bulb slices: GABA- and other transmitter receptors. J. Comp. Physiol. [A] 170, 153–159.

Burrows, M. (1989). Processing of mechanosensory signals in local reflex pathways of the locust. J. Exp. Biol. 146, 209–227.

Chou, Y.-H., Spletter, M.L., Yaksi, E., Leong, J.C.S., Wilson, R.I., and Luo, L. (2010). Diversity and wiring variability of olfactory local interneurons in the Drosophila antennal lobe. Nat Neurosci.

Christensen, T.A., Waldrop, B.R., Harrow, I.D., and Hildebrand, J.G. (1993). Local interneurons and information processing in the olfactory glomeruli of the moth Manduca sexta. J Comp Physiol A 173, 385–399.

Christensen, T.A., Waldrop, B.R., and Hildebrand, J.G. (1998). GABAergic mechanisms that shape the temporal response to odors in moth olfactory projection neurons. Ann N Acad Sci 855, 475–481.

Das, A., Chiang, A., Davla, S., Priya, R., Reichert, H., Vijayraghavan, K., and Rodrigues, V. (2011). Identification and analysis of a glutamatergic local interneuron lineage in the adult Drosophila olfactory system. Neural Syst Circuits 1, 4.

Das, S., Trona, F., Khallaf, M.A., Schuh, E., Knaden, M., Hansson, B.S., and Sachse, S. (2017). Electrical synapses mediate synergism between pheromone and food odors in Drosophila melanogaster. Proc. Natl. Acad. Sci. U. S. A. 114, E9962–E9971.

Demmer, H., and Kloppenburg, P. (2009). Intrinsic Membrane Properties and Inhibitory Synaptic Input of Kenyon Cells as Mechanisms for Sparse Coding? J. Neurophysiol. 102, 1538–1550.

Diamond, J.S. (2017). Inhibitory Interneurons in the Retina: Types, Circuitry, and Function. Annu. Rev. Vis. Sci. 3, 1–24.

Distler, P. (1989). Histochemical demonstration of GABA-like immunoreactivity in cobalt labeled neuron individuals in the insect olfactory pathway. Histochemistry 91, 245–249.

Distler, P. (1990). Synaptic connections of dopamine-immunoreactive neurons in the antennal lobes of Periplaneta americana. Colocalization with GABA-like immunoreactivity. Histochemistry 93, 401–408.

Distler, P.G., and Boeckh, J. (1997a). Synaptic connections between identified neuron types in the antennal lobe glomeruli of the cockroach, Periplaneta americana: I. Uniglomerular projection neurons. J Comp Neurol 378, 307–319.

Distler, P.G., and Boeckh, J. (1997b). Synaptik connections between identified neuron types in the antennal lobe glomeruli of the cockroach Periplaneta americana: II multiglomerular interneurons. J Comp Neurol 383, 529–540.

Distler, P.G., and Boeckh, J. (1997c). Synaptik connections between identified neuron types in the antennal lobe glomeruli of the cockroach Periplaneta americana: II multiglomerular interneurons. J Comp Neurol 383, 529–540.

Distler, P.G., Gruber, C., and Boeckh, J. (1998). Synaptic connections between GABA-immunoreactive neurons and uniglomerular projection neurons within the antennal lobe of the cockroach, Periplaneta americana. Synapse 29, 1–13.

Dodt, H.U., and Zieglgansberger, W. (1994). Infrared videomicroscopy: a new look at neuronal structure and function. Trends Neurosci. 17, 453–458.

Ennis, M., Puche, A.C., Holy, T., and Shipley, M.T. (2015). Chapter 27 - The Olfactory System. In The Rat Nervous System (Fourth Edition), G. Paxinos, ed. (San Diego: Academic Press), pp. 761–803.

Ernst, K.D., and Boeckh, J. (1983). A neuroanatomical study on the organization of the central antennal pathways in insects. III. Neuroanatomical characterization of physiologically defined response types of deutocerebral neurons in Periplaneta americana. Cell Tissue Res 229, 1–22.

Fujimura, K., Yokohari, F., and Tateda, H. (1991). Classification of antennal olfactory receptors of the cockroach, Periplaneta americana. Zool Sci 8, 243–255.

Fujiwara, T., Kazawa, T., Haupt, S.S., and Kanzaki, R. (2014). Postsynaptic odorant concentration dependent inhibition controls temporal properties of spike responses of projection neurons in the moth antennal lobe. PLoS One 9, e89132.

Fusca, D., Husch, A., Baumann, A., and Kloppenburg, P. (2013). Choline acetyltransferase-like immunoreactivity in a physiologically distinct subtype of olfactory nonspiking local interneurons in the cockroach (Periplaneta americana). J. Comp. Neurol. 521, 3556–3569.

Fusca, D., Schachtner, J., and Kloppenburg, P. (2015). Colocalization of allatotropin and tachykinin-related peptides with classical transmitters in physiologically distinct subtypes of olfactory local interneurons in the cockroach (Periplaneta americana). J Comp Neurol 523, 1569–1586.

Galizia, C.G., and Kimmerle, B. (2004). Physiological and morphological characterization of honeybee olfactory neurons combining electrophysiology, calcium imaging and confocal microscopy. J. Comp. Physiol. A Neuroethol. Sens. Neural. Behav. Physiol. 190, 21–38.

Hamill, O.P., Marty, A., Neher, E., Sakmann, B., and Sigworth, F.J. (1981). Improved patch-clamp techniques for high-resolution current recording from cells and cell-free membrane patches. Pflugers Arch 391, 85–100.

Hoskins, S.G., Homberg, U., Kingan, T.G., Christensen, T.A., and Hildebrand, J.G. (1986). Immunocytochemistry of GABA in the antennal lobes of the sphinx moth Manduca sexta. Cell Tissue Res 244, 243–252.

Huang, J., Zhang, W., Qiao, W., Hu, A., and Wang, Z. (2010). Functional connectivity and selective odor responses of excitatory local interneurons in Drosophila antennal lobe. Neuron 67, 1021–1033.

Husch, A., Paehler, M., Fusca, D., Paeger, L., and Kloppenburg, P. (2009a). Calcium current diversity in physiologically different local interneuron types of the antennal lobe. J Neurosci 29, 716–726.

Husch, A., Paehler, M., Fusca, D., Paeger, L., and Kloppenburg, P. (2009b). Distinct electrophysiological properties in subtypes of nonspiking olfactory local interneurons correlate with their cell type-specific Ca2+ current profiles. J Neurophysiol 102, 2834–2845.

Klagges, B.R., Heimbeck, G., Godenschwege, T.A., Hofbauer, A., Pflugfelder, G.O., Reifegerste, R., Reisch, D., Schaupp, M., Buchner, S., and Buchner, E. (1996). Invertebrate synapsins: a single gene codes for several isoforms in Drosophila. J. Neurosci. 16, 3154–3165.

Kloppenburg, P., Ferns, D., and Mercer, A.R. (1999). Serotonin enhances central olfactory neuron responses to female sex pheromone in the male sphinx moth manduca sexta. J Neurosci 19, 8172–8181.

Laurent, G., and Davidowitz, H. (1994). Encoding of olfactory information with oscillating neural assemblies. Science 265, 1872–1875.

Lemon, and Getz (1997). Temporal resolution of general odor pulses by olfactory sensory neurons in American cockroaches. J. Exp. Biol. 200, 1809–1819.

Lemon, W.C., and Getz, W.M. (1998). Responses of cockroach antennal lobe projection neurons to pulsatile olfactory stimuli. Ann. N. Y. Acad. Sci. 855, 517–520.

Lemon, W.C., and Getz, W.M. (2000). Rate code input produces temporal code output from cockroach antennal lobes. Biosystems 58, 151–158.

Liu, W.W., and Wilson, R.I. (2013). Glutamate is an inhibitory neurotransmitter in the Drosophila olfactory system. Proc Natl Acad Sci U A 110, 10294–10299.

Liu, S., Puche, A.C., and Shipley, M.T. (2016). The Interglomerular Circuit Potently Inhibits Olfactory Bulb Output Neurons by Both Direct and Indirect Pathways. J. Neurosci. Off. J. Soc. Neurosci. 36, 9604–9617.

MacLeod, K., and Laurent, G. (1996). Distinct mechanisms for synchronization and temporal patterning of odor-encoding neural assemblies. Science 274, 976–979.

Malun, D. (1991a). Inventory and distribution of synapses of identified uniglomerular projection neurons in the antennal lobe of Periplaneta americana. J Comp Neurol 305, 348– 360.

Malun, D. (1991b). Synaptic relationships between GABA-immunoreactive neurons and an identified uniglomerular projection neuron in the antennal lobe of Periplaneta americana: a double-labeling electron microscopic study. Histochemistry 96, 197–207.

Malun, D., Waldow, U., Kraus, D., and Boeckh, J. (1993). Connections between the deutocerebrum and the protocerebrum, and neuroanatomy of several classes of deutocerebral projection neurons in the brain of male Periplaneta americana. J Comp Neurol 329, 143–162.

Mohamed, A.A., Retzke, T., Chakraborty, S.D., Fabian, B., Hansson, B.S., Knaden, M., and Sachse, S. (2019). Odor mixtures of opposing valence unveil inter-glomerular crosstalk in the Drosophila antennal lobe. Nat. Commun. 10, 1201.

Nagel, K.I., and Wilson, R.I. (2016). Mechanisms Underlying Population Response Dynamics in Inhibitory Interneurons of the Drosophila Antennal Lobe. J. Neurosci. 36, 4325–4338.

Najac, M., Diez, A.S., Kumar, A., Benito, N., Charpak, S., and De Saint Jan, D. (2015). Intraglomerular lateral inhibition promotes spike timing variability in principal neurons of the olfactory bulb. J. Neurosci. 35, 4319–4331.

Neupert, S., Fusca, D., Schachtner, J., Peter Kloppenburg, and Predel, R. (2012). Toward a single-cell-based analysis of neuropeptide expression in Periplaneta americana antennal lobe neurons. J Comp Neurol 520, 694–716.

Neupert, S., Fusca, D., Kloppenburg, P., and Predel, R. (2018). Analysis of single neurons by perforated patch clamp recordings and MALDI-TOF mass spectrometry. ACS Chem. Neurosci. 9, 2089–2096.

Nishino, H., Iwasaki, M., Kamimura, I., and Mizunami, M. (2012). Divergent and convergent projections to the two parallel olfactory centers from two neighboring, pheromone-receptive glomeruli in the male American cockroach. J Comp Neurol 520, 3428–3445.

Nishino, H., Watanabe, H., Kamimura, I., Yokohari, F., and Mizunami, M. (2015). Coarse topographic organization of pheromone-sensitive afferents from different antennal surfaces in the American cockroach. Neurosci Lett 595, 35–40.

Nishino, H., Iwasaki, M., Paoli, M., Kamimura, I., Yoritsune, A., and Mizunami, M. (2018). Spatial Receptive Fields for Odor Localization. Curr. Biol. CB 28, 600-608.e3.

Oliveira, E.E., Pippow, A., Salgado, V.L., B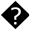 schges, A., Schmidt, J., and Kloppenburg, P. (2010). Cholinergic currents in leg motoneurons of Carausius morosus. J Neurophysiol 103, 2770–2782.

Olsen, S.R., and Wilson, R.I. (2008). Lateral presynaptic inhibition mediates gain control in an olfactory circuit. Nature 452, 956–960.

Olsen, S.R., Bhandawat, V., and Wilson, R.I. (2007). Excitatory interactions between olfactory processing channels in the Drosophila antennal lobe. Neuron 54, 89–103.

Paeger, L., Bardos, V., and Kloppenburg, P. (2017). Transient voltage-activated K(+) currents in central antennal lobe neurons: cell type-specific functional properties. J. Neurophysiol. 117, 2053–2064.

Paoli, M., Nishino, H., Couzin-Fuchs, E., and Galizia, C.G. (2020). Coding of odour and space in the hemimetabolous insect Periplaneta americana. J. Exp. Biol. 223.

Pearson, K.G., and Fourtner, C.R. (1975). Nonspiking interneurons in walking system of the cockroach. J. Neurophysiol. 38, 33–52.

Pippow, A., Husch, A., Pouzat, C., and Kloppenburg, P. (2009). Differences of Ca2+ handling properties in identified central olfactory neurons of the antennal lobe. CELL CALCIUM 46, 87–98.

Root, C.M., Semmelhack, J.L., Wong, A.M., Flores, J., and Wang, J.W. (2007). Propagation of olfactory information in Drosophila. Proc. Natl. Acad. Sci. U. S. A. 104, 11826–11831.

Sachse, S., and Galizia, C. (2002). Role of inhibition for temporal and spatial odor representation in olfactory output neurons: a calcium imaging study. J Neurophysiol 87, 1106– 1117.

Sachse, S., Rappert, A., and Galizia, C.G. (1999). The spatial representation of chemical structures in the antennal lobe of honeybees: steps towards the olfactory code. Eur. J. Neurosci. 11, 3970–3982.

Sattelle, D. (1992). Receptors for L-glutamate and GABA in the nervous system of an insect (Periplaneta americana). Comp. Biochem. Physiol. Part C Comp. Pharmacol. 103, 429–438.

Seki, Y., Rybak, J., Wicher, D., Sachse, S., and Hansson, B.S. (2010). Physiological and morphological characterization of local interneurons in the Drosophila antennal lobe. J Neurophysiol 104, 1007–1019.

Shang, Y., Claridge-Chang, A., Sjulson, L., Pypaert, M., and Miesenbock, G. (2007). Excitatory Local Circuits and Their Implications for Olfactory Processing in the Fly Antennal Lobe. Cell 128, 601–612.

Shepherd, G.M., Chen, W.R., and Greer, C.A. (2004). Olfactory Bulb. In The Synaptic Organization of the Brain, (Oxford University Press), p.

Silbering, A.F., Rytz, R., Grosjean, Y., Abuin, L., Ramdya, P., Jefferis, G.S.X.E., and Benton, R. (2011). Complementary function and integrated wiring of the evolutionarily distinct Drosophila olfactory subsystems. J. Neurosci. Off. J. Soc. Neurosci. 31, 13357–13375.

Strausfeld, N.J., and Li, Y.S. (1999). Representation of the calyces in the medial and vertical lobes of cockroach mushroom bodies. J Comp Neurol 409, 626–646.

Tabuchi, M., Dong, L., Inoue, S., Namiki, S., Sakurai, T., Nakatani, K., and Kanzaki, R. (2015). Two types of local interneurons are distinguished by morphology, intrinsic membrane properties, and functional connectivity in the moth antennal lobe. J Neurophysiol jn.00050.2015.

Wachowiak, M., and Shipley, M.T. (2006). Coding and synaptic processing of sensory information in the glomerular layer of the olfactory bulb. Semin. Cell Dev. Biol. 17, 411–423.

Waldrop, B., Christensen, T.A., and Hildebrand, J.G. (1987). GABA-mediated synaptic inhibition of projection neurons in the antennal lobes of the sphinx moth, Manduca sexta. J Comp Physiol A 161, 23–32.

Warren, B., and Kloppenburg, P. (2014). Rapid and Slow Chemical Synaptic Interactions of Cholinergic Projection Neurons and GABAergic Local Interneurons in the Insect Antennal Lobe. J. Neurosci. 34, 13039–13046.

Watanabe, H., Nishino, H., Nishikawa, M., Mizunami, M., and Yokohari, F. (2010). Complete mapping of glomeruli based on sensory nerve branching pattern in the primary olfactory center of the cockroach Periplaneta americana. J Comp Neurol 518, 3907–3930.

Watanabe, H., Haupt, S.S., Nishino, H., Nishikawa, M., and Yokohari, F. (2012). Sensillum-specific, topographic projection patterns of olfactory receptor neurons in the antennal lobe of the cockroach Periplaneta americana. J Comp Neurol 520, 1687–1701.

Watanabe, H., Nishino, H., Mizunami, M., and Yokohari, F. (2017). Two Parallel Olfactory Pathways for Processing General Odors in a Cockroach. Front. Neural Circuits 11, 32.

Wellis, D.P., and Scott, J.W. (1990). Intracellular responses of identified rat olfactory bulb interneurons to electrical and odor stimulation. J. Neurophysiol. 64, 932–947.

Wilson, R.I. (2013). Early olfactory processing in Drosophila: mechanisms and principles. Annu Rev Neurosci 36, 217–241.

Wilson, R.I., and Laurent, G. (2005). Role of GABAergic inhibition in shaping odor-evoked spatiotemporal patterns in the Drosophila antennal lobe. J Neurosci 25, 9069–9079.

Yaksi, E., and Wilson, R.I. (2010). Electrical coupling between olfactory glomeruli. Neuron 67, 1034–1047.

